# The Conquering influence of the Nano extracts of pomegranate peels and pistachio leaves in amelioration of acute liver failure

**DOI:** 10.1101/2020.05.06.071001

**Authors:** Sohair Aly Hassan, Olfat Hammam, Sahar Awdallah Hussein, Wessam Magdi Aziz

## Abstract

**Objective:** Acute liver failure (ALF) is a clinical condition with an unclear history of pathophysiology, making it a challenging task for scientists to reverse the disease in its initial phase and to help the liver re-function as usual: this study was proposed to estimate the hepatoprotective effects of Punicagranatum peel and ***Pistaciaatlantica*** leaves as a multi-rich antioxidants ingredients either in their normal and/or in their nano forms against thioacetamide induced acute liver failure in a rodent model.

**Method:** Sixty male Wistar rats were divided into six equal groups, the first group employed as a control; the second group was given a dose of thioacetamide (TAA)-350 mg/bwip., from the third to the sixth group received TAA □ 2ml of 100gm of aqueous extracts of ***PunicagranatumL*** and ***Pistaciaatlantica*** either in their normal and/or Nano forms consecutively along 2 weeks.

**Results:** recorded significant elevation in liver enzymes, lipid profiles, LPO and NO with marked significant decrease in GSH and SOD accompanied by elevation in inflammatory cytokine (IL6,TNF-α and AFP) in addition to a noticeable increase in HSP_70_ level & degradation in DNA respectively in TAA challenged group. However significant and subsequent amelioration of most of the impaired markers has been observed with ipnano treatment of both extracts.

**Conclusion:** The current results highlighted the high performance of both plant nano extracts and their hepatoprotective impact and their possible therapeutic role in the amelioration of TAA induced acute liver failure in experimental animals.

## 1. Introduction

Acute failure of the liver/ALF is a disease characterized by a rapid and significant impairment of hepatocyte function in patients with normal liver function [1]. Hepatic encephalopathy and death levels of up to 80% are caused by ALF, even if intensive care is given [2,3,4,5,6]. Liver transplantation has been shown to be the most effective therapy for ALF, but this therapeutic choice constrained by the lak of donor organ. The use of bio artificial livers is one of the other approaches to treatment, to overcome the time gap for transplantation or regeneration of the liver [7,4,8,9]. The effectiveness of new ALF treatment methods has not yet been fully explored [5, 9] and even the knowledge of the ALF pathophysiology is still inadequate [2, 6]. So there is a necessitates to search for the discovery and implementation of efficacious safe nutraceuticals rather than chemically synthesized as well as an Optimum experimental ALF models with features similar to human disorder.

Thioacetamide (TAA) is a class of chemicals that most recommended to generate the disease model in the very shortest duration^[10, 11]^. TAA has long been known as a hepatocyte toxic agent; The TAA biotransformation cycle into thioacetamide sulfoxide (TAAS) mainly takes place through P450 (CYP) cytochrome pathway, (CYP2E enzyme is mainly involved here)^[12]^. The binding of these metabolites to liver macromolecules dramatically inspire a moiety of reactive oxygen species in turn induces acute centrilobular necrosis of the liver^[13]^. These results of the liver diseases such as cirrhosis and primary cancer of the liver^[14]^ TAA will overpower the body’s antioxidant defense mechanisms and spread rapidly to neighboring tissues to destroy cell vital ingredients that ultimately enter DNA.^[15]^. There is currently a widespread interest in the use of natural products specially the dietary polyphenols, which are considered safe and less side-effect compared to the synthesis of chemical drugs and the best hepatoprotective agents due to their synergistic action^[16, 17]^. Dietary polyphenols, especially those in vegetables, fruits and legumes, have been reported to be capable of achieving a balance between natural antioxidants and free radicals by enhancing the role of natural antioxidant enzymes such as glutathione reductase (GR superoxide dismutase (SOD), glutathione peroxidase (GPx), and glutathione S transferase (GST) by specifically capturing free radicals^[18]^.

Sweet fruit belongs to *Punicaceae* family is known worldwide as pomegranate / ***Punicagranatum L*** because of its health value from all its components^[19,20, 23]^. Although anti-inflammatory properties and resistance to exponential increases in glucose levels in diabetic rat models have been reported for pomegranate flowers and juices^[21,22]^More and more attention has been paid recently to the pomegranate husk / peel as it has been confirmed to be of great benefit to a broad spectrum of hepatic inflammation and other liver-related diseases^[24,25]^. This conclusion assigned according to earlier studies has shown that pomegranate peels have a higher content of phenolic compounds than fruit pulp including a group of flavonoids (catechins, anthocyanins and other complex flavonoids) and hydrolyzing tannins (punicalin, pedunculagin, punicalagin, gallic and ellagic acid)^[21,26]^. Pomegranate has shown evidence concerning preventive cancer-chemo property^[27]^. Reducing the oxidative stress mediators that extensively contribute to its phenolic compounds and serve as a powerful tool for free radical scavenging properties makes it unique^[28]^. Pistachio is a type of nut produced by approximately 20 species of shrub such as ***Pistaciaterebinthusn, Pistaciavera, Pistaciakhinjuk, Pistacialentiscus***, and ***Pistaciaatlantica*** belonging to the Cashew family (***Anacardiaceae***)^[29]^.This Genus Pistacia is characterized by the presence of a wide range of diverse compounds such as flavonoids, triterpenes, sterols and phenolic compounds. The phenolic compounds are abundant in different parts of this genus^[30,31]^. ***Pistaciaatlantica*** has been shown to have antibacterial properties and is used in eczema, throat infections, kidney stones, asthma and stomach aches as an astringent, antipyretic, anti-inflammatory, antiviral, and stimulating agent^[32]^. Previous studies of Photochemistry and chromatographic fingerprints of this plant indicated that it contain phenolic compounds (such as gallic acid, quinic acid, tetragall oylquinic acid,) and triterpenoids, pinene, terpinolene^[33]^. Although the pistachio is common, it was not sufficiently investigated as a safe eaten nut^[34]^. The use of nanoparticles science is completely exhibit an improved novel properties compared with larger particles of the bulk materials due to the variations in the particles size, morphology and distribution. Nowadays, noble metals like silver, gold and platinum are commonly used in green chemistry without using toxic chemicals via a method involve use of environmentally friendly processes as a simple and viable alternative to synthetic chemicals and physical processes^[35]^. Several researchers reported the biosynthesis and possible applications of metal nanoparticles by plant leaf extracts^[36]^.

This work is the first to clarify the potential role of pomegranate peel and pistachio leaves extracts in their natural and nano forms to alleviate TAA-induced hepatic failure in rat model. This research further explores a potential mechanism of action for both forms that could have therapeutic effects.

## 2. Materials and method

### 2.1 Chemicals

All Chemicals, Silver nitrate and thioacetamide were procured from Sigma-Aldrich (USA). The different solvents and chemicals used in the present study were of high analytical grade and purchased from a local vendor.

### 2.2 Authentication & preparation of the plant extracts

The pomegranate fruits were purchased from local market at Cairo city, Egypt, 2018 and ***Pistaciaatlantica*** leaves were collected from Beniwalid, Libya 2017.The plants material were identified and authenticated by a plant taxonomist at the National Research Center Herbarium (Cairo, Egypt) according to ^[37]^. Where a voucher specimen has been deposited for future reference. The pomegranates was washed, cut into small pieces and left to dry in air and the complete dryness occurred in an oven at 50°C. The dried peels were grounded to fine powder. The pomegranate peel (PO) extract was obtained by addition 100 gm of pomegranate peel powder with 1000 ml boiled distilled water [Erlenmeyer flask] and boiled for 10 minutes with the help of whatman filter paper(NO.3), the boiled materials were filtered toget aqueous fruit peel extract,which was used as such for metal nanoparticles synthesis. Finally stored at −4 °C for further experiments. About 100 g of dried powder of ***Pistaciaatlantica*** leaves (PS) transferred into a 1000-mL Erlenmeyer flask with 500 mL of distilled water along with boiling for 30 min. The aqueous extract was filtered and stored at 4 °C^[33]^.

#### High performance liquid Chromatography [HPLC]

HPLC analysis was carried out using an Agilent 1260 series. The separation was carried out using Kromasil C18 column (4.6 mm x 250 mm i.d., 5 μm).The mobile phase consisted of water (A) and 0.05% trifluoroacetic acid in acetonitrile (B) at a flow rate 1 ml/min. The mobile phase was programmed consecutively in a linear gradient as follows: 0 min (82% A); 0 –5 min (80% A); 5-8 min (60% A); 8-12 min (60% A); 12-15 min (85% A) and 15-16 min (82% A). The multi-wavelength detector was monitored at 280 nm. The injection volume was 10 μl for each of the sample solutions. The column temperature was maintained at 35 °C. [**Chart 1a,b]**,while ***Pistaciaatlantica*** was evaluated according to the methods described for the identification of major phytochemical components in the extract^[38]^.

### 2.3. Synthesis of nanoparticles

To synthesis silver nanoparticle (1MLAgNPs), silver nitrate (0.017 g) was added in 100 ml of double distilled water to prepare 1 mM silver nitrate solution. Then, 1 ml of aqueous extract of both extracts was added to 50 ml of silver nitrate (1 mM) solution to obtain silver nanoparticles, as shown in Fig1a, b.

**Fig. 1:**
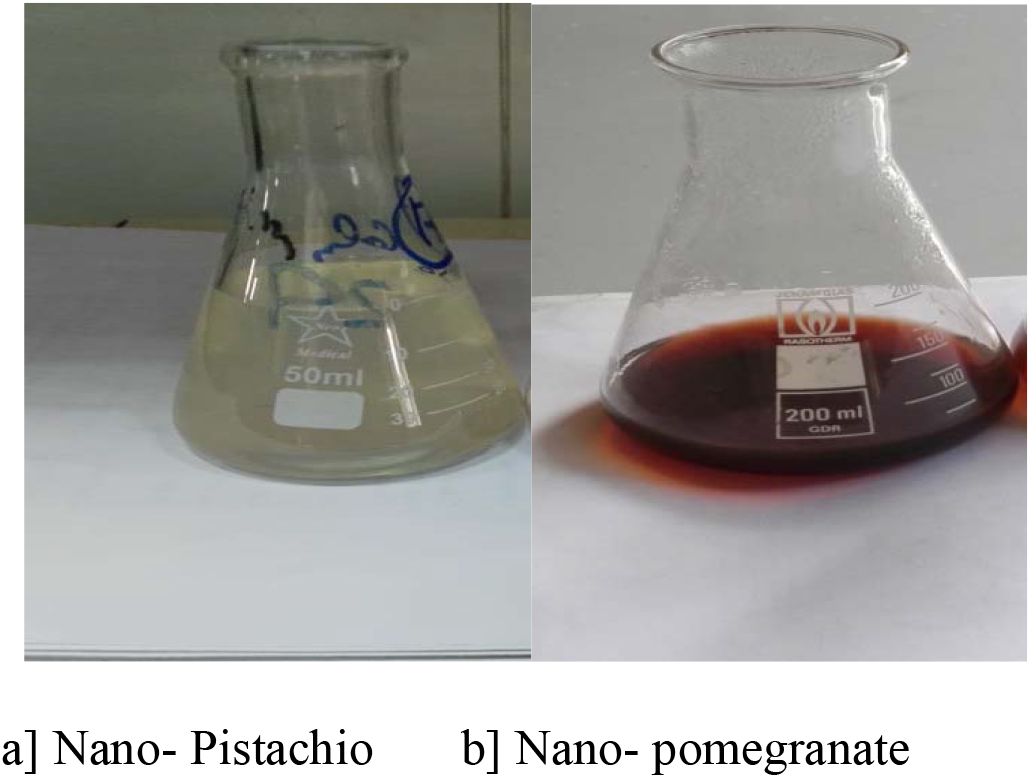
The visual observation of color changes of nano extracts.

This setup was incubated in a dark chamber to minimize photo-activation of silver nitrate at room temperature under static conditions for nucleation of the silver nanoparticles. Reduction of Ag to AgO^2^ was confirmed by the color change of solution from colorless to brown^[39]^. The suspensions were subjected to centrifugation for the pellets to settle, which were then dried using a vacuum drier18 2.5 Characterization of the silver nanoparticles to measure the maximum production and activity of AgNPs, UV-visible spectra were taken using a Shimadzu UV-visible spectrophotometer.

### 2.4. Characterization of synthesized silver nanoparticles

#### 2.4.1. UV-Vis spectral analysis

was done by using Shimadzu Visible spectrophotometer (UV-1800, Japan). The periodic scans of the optical absorbance between 300 and 700 nm with a double-beam UV-Visible spectrophotometer (Carry 100 with tungsten halogen light sources) were performed to investigate the reduction of silver ions by leaf extract. The presented results were obtained at room temperature.

#### 2.4.2. Field emission scanning electron microscopy (FESEM)

The morphology, particle dispersion, and chemical composition of the prepared Nano-structures were investigated by (FESEM) (Quanta 450) equipped with EDS at accelerating voltage 30 kV. Cross cut samples were prepared by fracturing the membranes in liquid nitrogen. A thin layer of gold was coated on all the samples before microscopic analysis

#### 2.4.3. The potential of the biosynthesized AgNPs

was performed on Malvern Zeta size Nano series compact scattering spectrometer (Malvern Instruments Ltd.; Malvern, UK).

#### 2.4.4. Transmission electron microscopy (TEM)

The particle size and surface morphology were analyzed usinga JEOL JEM 1230 transmission electron microscope (JEOL Ltd., USA). The samples for TEM analysis were cut into slices of a nominal thickness of 100 nm using an ultra-microtome with a diamond knife at an ambient temperature. The cut samples were supported on a copper mesh for this analysis.

### 2.5. Animals

A total of 60 Wistar ratsaged 2 months weighing 180–200 g, were taken from the breeding laboratory of National research center and kept in animal care facility. Rats were kept in polypropylene cages at room temperature with a 12 h light-dark cycle and relative humidity of 60% ± 1 %. Animals were provided purified water ad libitum with standard laboratory rat chow. The study was approved by the animal ethics committee of Institutional Review Board of National Research Center, and the ethical guidelines for use and care of laboratory animals in experiments were strictly followed.The experimental animal protocol was designed to minimize pain or discomfort to the animals. TAA (Sigma, Aldrich Chemical Co., USA)) was dissolved in physiological saline and the appropriate selected dose was injected 350 mg/BW i.p. in 1-ml volume^[40]^.

#### 2.5.1. Experimental protocol

The rats were divided into 6 groups consisting of 10 rats each. Rats injected intraperitoneal (IP) with freshly prepared TAA 350 gm / BW a single injection^[40]^. 5 % glucose solution was added to the drinking water to prevent dehydration 24 hrs post injection, daily IP dose of 2mls of normal & Nano pomegranate extracts, were given to the third and fourth group respectively. The fifth & sixth groups were given the normal and nano forms of pistachio extracts in a dose of 2mls along two weeks. While the first group and second groups were kept as negative and positive control groups. The survival rate & BW was monitored along the experiment period. 24 hr after the last dose of respective assigned treatments, the whole animals were sacrificed by cervical decapitation. Blood samples were immediately collected into sterilized tubes and let stand for 10 minutes to clot. The serum was separated by centrifugation at 3000 rpm for 10 min and stored at-80° C for assessment of liver functions and another biochemical analysis. Liver tissues were dissected and divided into two parts, the first part was fixed in 10% formaldehyde solution for histopathological and immunohistochemical examinations and the second part (1 g) was homogenized (1:5 w/v) in ice cold 0.1 M potassium phosphate buffer (pH 7.4). The homogenate was centrifuged at 4,000 rpm for 15 minutes and the supernatant collected and liquated in eppendorf tubes and stored a −20 °C for different biochemical determinations.

### 2.6. Biochemical analysis

The liver function indices, Serum alanine aminotransferase (ALT), aspartate aminotransferase (AST) and alkaline phosphatase (ALP),total lipid profile (TP), total cholesterol (TC), triglyceride (TG) and total bilirubin were estimatedusing biodiagnostic kits (Biogamma, Stanbio, WestGermany). The oxidative stress markers, lipid peroxidation (LPO), glutathione (GSH), superoxide dismutase (SOD) and nitric oxide (NO), were evaluated using the methods of ^[40,41,42,43, 44]^ respectively. Liver total protein contents were estimated by the method of ^[45]^. The DNA fragmentation index was evaluated using the comet assay technique as described by^[46]^. The proinflammatory cytokines TNF-α, IL-6, Alpha fetoprotein (AFP) and HSP 70 were assayed using commercially available ELISA kits (R&D, Minneapolis, MN).

### 2.7. Histopathological studies

A portion of the liver was preserved in 10% buffer formalin for at least 24 hr. and then processed and embedded in paraffin according to the standard protocol. Sections were cut into 5 μm thick portions, transferred on to glass slides, and stained with haematoxylin and eosin (HE), observed microscopically for any histopathological and immunohistochemical changes as well^[47]^.

#### 2.7.1. Histopathological and immunohistochemical examinations

Five μm thick sections of paraffinized liver stained with hematoxylin and eosin were examined histopathologically under Zeiss microscope (Carl Zeiss Microscopy GmbH 07745 Jena, Germany) with × 200 magnification powers. For immunohistochemical examinations of IL6, TNF α were performed on liver sections cuts from the paraffin blocks with a commercially available Anti-mouse IL6, TNF α antibodies (Santa Cruz Biotechnology, CA, USA) at the optimal working dilution of 1:100. Briefly, slides were sectioned at 4 μm on to positively charged slides (Superfrost plus, Menzel-Glaser, Germany) and the slides were stained on an automated platform (DakoAutostainer Link 48). Heat induced antigen retrieval was used for 30 min at 97°C in the high-PH EnVision™ FLEX Target Retrieval Solution and the primary antibody was used at a dilution of 1 in 100.The percent of positively stained brown cytoplasm (IL6, TNF α) were examined in 10 microscopic fields (under Zeiss light microscopy at x400).

### 2.8. Statistical analysis

All data were expressed as mean ± SD of ten rats in each group using the Co-stat computer program where an unshared letter is significant at *p*□<□0.05. The correlations between all markers and inflammatory cytokines were analyzed using Pearson’s correlation analysis test. The ROC-Curve method was done by SPSS version 16 and used to generate the Cut-off value. area under the curve, Sensitivity, Specificity. P < 0.05 was set to be level of significance^[48]^.

## 3. Results

The current results revealed that the incubation of 50 ml of 1 mM AgNO_3_ with 1 ml of pomegranate extract for 24 h at the room temperature led to the synthesis of AgNPs as indicated by the development of the brown color (**Fig.1**).

Results of HPLC for pomegranate showed Gallic acid, chlorogenic acid and ellagic acid at concentrations of 2160.89, 763.22 and 97.60 were the major components of the extract (Table 1, chart 1a,b) while twenty-seven polyphenol compounds were detected in pistachio^[38]^, consisting of three major polyphenol groups, namely gallate, flavonoid and ellagic acid derivatives.

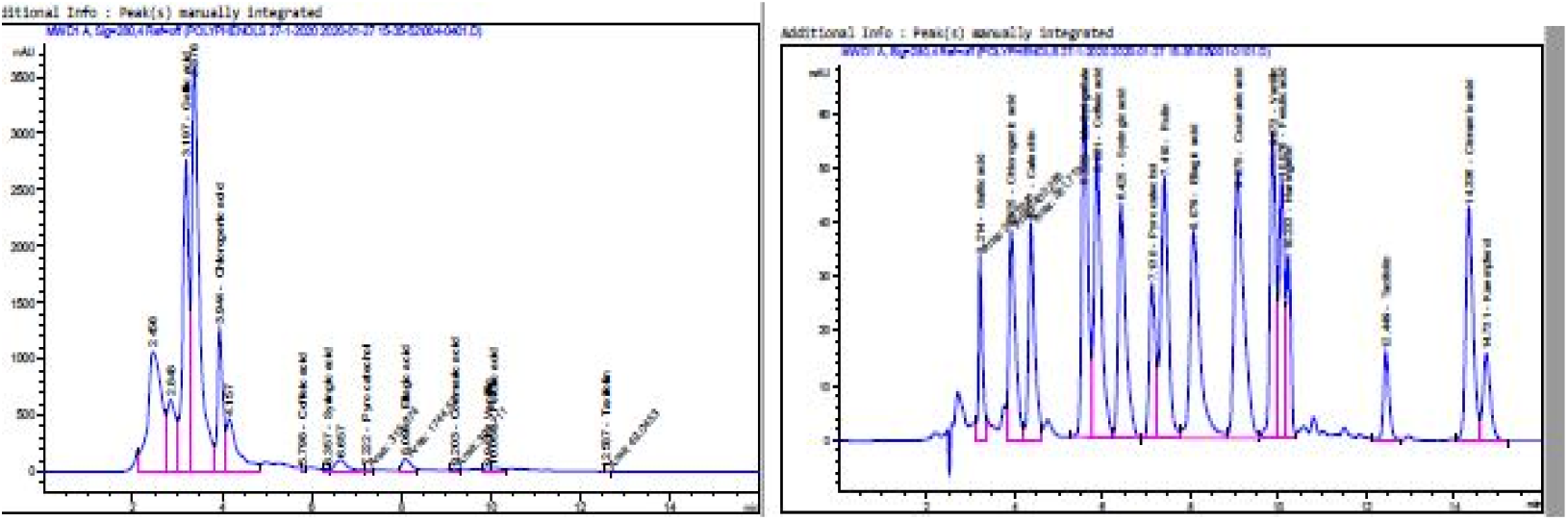
**Chart [1a]**:.High-performance liquid chromatography (HPLC) of polyphenolic compounds detected in water extracts of pomegranate peel water extract **Chart [1b]**: High-performance liquid chromatography (HPLC) of Standared polyphenolic compounds.

**Table 1:** High-performance liquid chromatography of polyphenolic compounds detected in aqueous extracts of ***Punica granatum L***

| CONTENTS | RTIME<br>[MIN] | CONC. ($\mu\text{G}/\text{ML} = \mu\text{G}/0.1\text{ G}$) |
| --- | --- | --- |
| Gallic acid | 3.214 | 2160.89 |
| Chlorogenic acid | 3.925 | 763.22 |
| Catechin | 4.377 | 0.00 |
| Methyl gallate | 5.589 | 0.00 |
| Coffeic acid | 5.881 | 8.05 |
| Syringic acid | 6.425 | 8.22 |
| Pyro catechol | 7.130 | 32.00 |
| Rutin | 7.410 | 0.00 |
| Ellagic acid | 8.079 | 97.60 |
| Coumaric acid | 9.075 | 6.98 |
| Vanillin | 9.873 | 9.55 |
| Ferulic acid | 10.076 | 18.78 |
| Naringenin | 10.233 | 0.00 |
| Taxifolin | 12.449 | 4.67 |
| Cinnamic acid | 14.336 | 0.00 |
| Kaempferol | 14.731 | 0.00 |

### 3.1. Characterization of silver nanoparticles by UV-VIS absorption spectroscopy

The absorption spectra of aqueous peel extracts obtained from Pomegranate plants, aqueous leaves extracts obtained from Pistachio were compared with the absorption spectra of silver nanoparticles prepared using these extracts in order to reveal the formation of silver phytonanoparticles. Further, UV–visible spectrophotometric analysis of nanoparticles showed the maximum absorption of nanoparticles of both PO and PS at a wavelength of 425 nm.

### 3.2. Electron Microscope scanning

Scanning electron microscope analysis showed that the synthesized AgNPs are polydispersed with particle size range from 19-65 nm for Pistachio leaves **Fig. 2.** and 12-18 nm for Pomegranate peel **Fig. 3**.

**Fig. 2.**
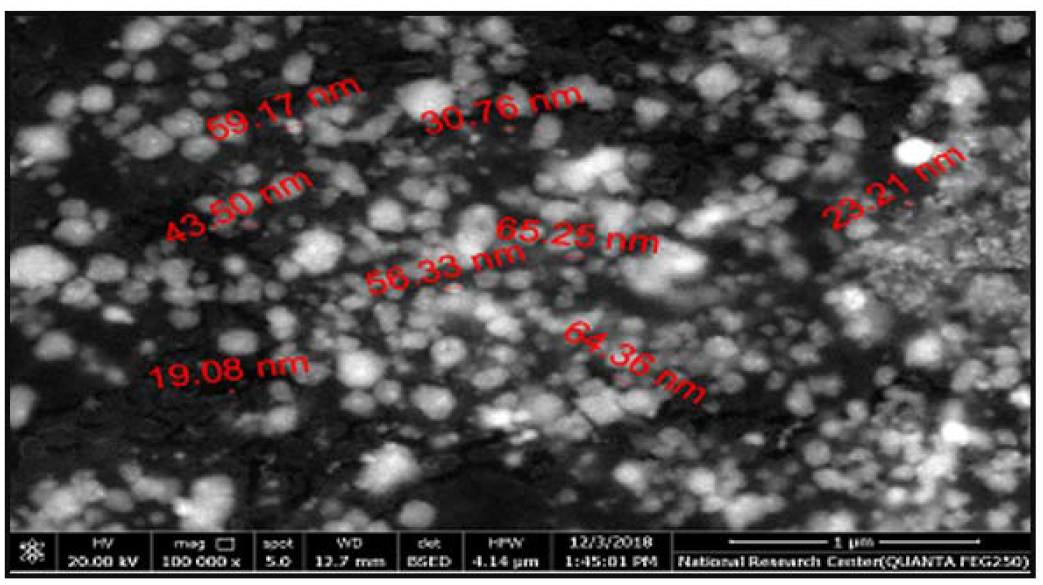
SEM Pistachio nanoparticle micrograph with a size range from 19-65 nm.

**Fig. 3.**
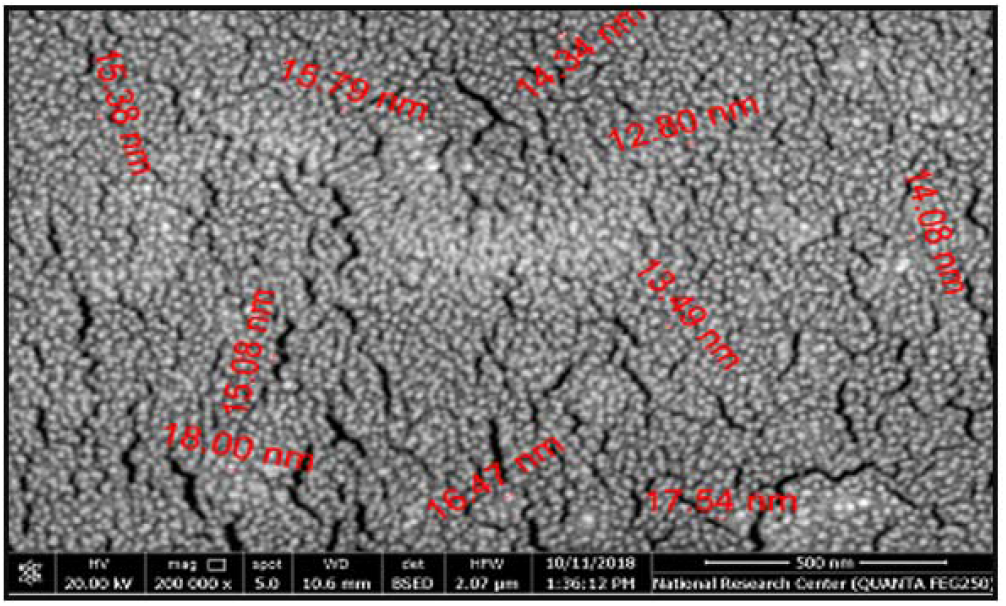
SEM Pomegranate **nanoparticle micrograph with a size range from 12-18 nm.**

### 3.3. Evaluation of physical stability of silver phytonanoparticles

The physical stability of the silver nanostructures was evaluated in terms of ξ-potential. The surface charge of all the silver nanostructures of Pomegranate peel and Pistachio leaves were ranging from −10.6 to −13.3 mV, −17.6-23.6 mV respectively.

### 3.4. Transmission electron microscopy (TEM)

TEM findings confirmed the spherical shape of the synthesized AgNPs with an average range of 4 to 42 nm, showing that the AgNPs pomegranate size is 17 nm Fig.4, whereas the AgNPs pistachio average size was 15 nm Fig 5.

**Fig. 4.**
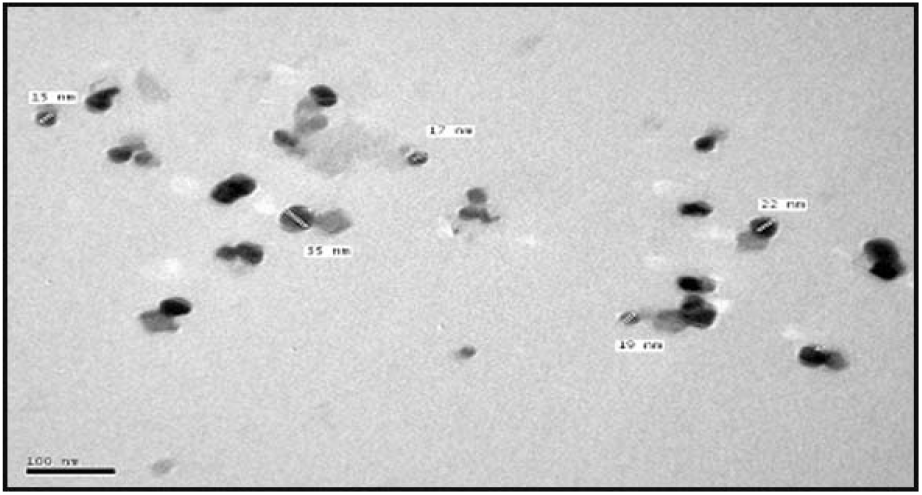
Pomegranate silver nanoparticles TEM micrograph, appear in spherical shape with mean size of 17 nm

**Fig. 5.**
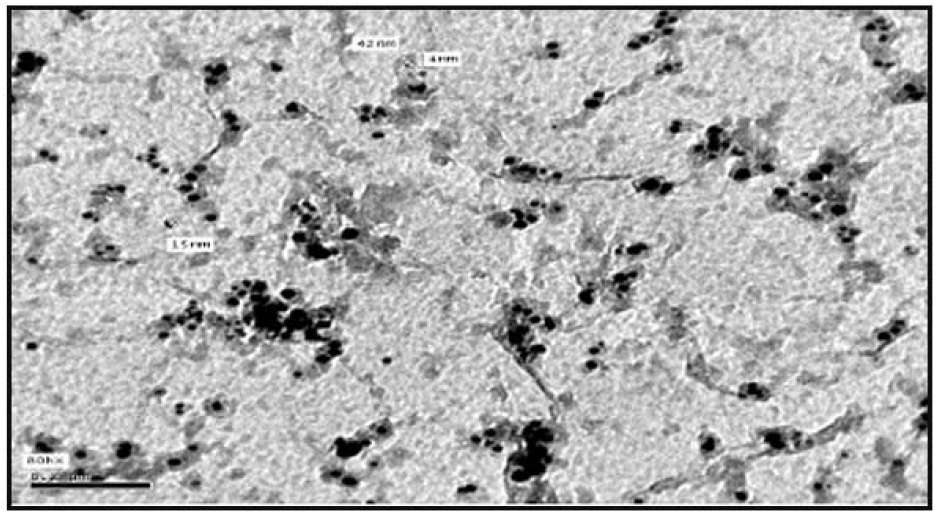
TEM micrograph of Pistachio silver nanoparticles, appear in a-spherical shapewith average mean size of 15 nm

### 3.5. Effect of pomegranate and pistachio in their normal and Nano forms on liver functions, lipid profile parameter

Upon administration of TAA, there was a marked increase in serum activity, liver function represented by (ALT, AST and ALP) as well as bilirubin to approximately 2.5, 2.8, 2.15 and 1.93 fold, respectively, as well as a marked increase in the lipid profile represented by TP, TG, TC to approximately 2.6, 3.2.2 fold, respectively, compared to the normal control group at ***p***<0.0001 **(Table 2)**. Treatment of animals with pomegranate peel (PO) and pistachio leaves (PS) extract either in their normal or in Nano forms (100 mg/kg) significantly decreased ALT, AST, ALP levels. A notable increase in the level of bilirubin in comparison with the challenged TAA group at ***p*** < 0.0001 **(Table 2)**. Hence, treatment showed very satisfied improvement levels as shown in **Fig. 6.** Nano pistachio group showed the most highly improved % in liver function tests. On the other hand lipid profiles upon treatment with PO and PS in normal and Nano form a prominent significant reduction in (TP, TC, and TG) were recorded compared with TAA group at ***p*** < 0.0001 **(Table 2).** Therefore, we observed amelioration levels in these parameters as represented in Fig. 7. Whereas in total lipids, Pomegranate extract obtained the most percent improvement other than the remaining treatment classes, while Pistachio leaves extract showed a high percentage improvement in TC and TG parameters. **(Fig. 7).**

**Fig. 6:**
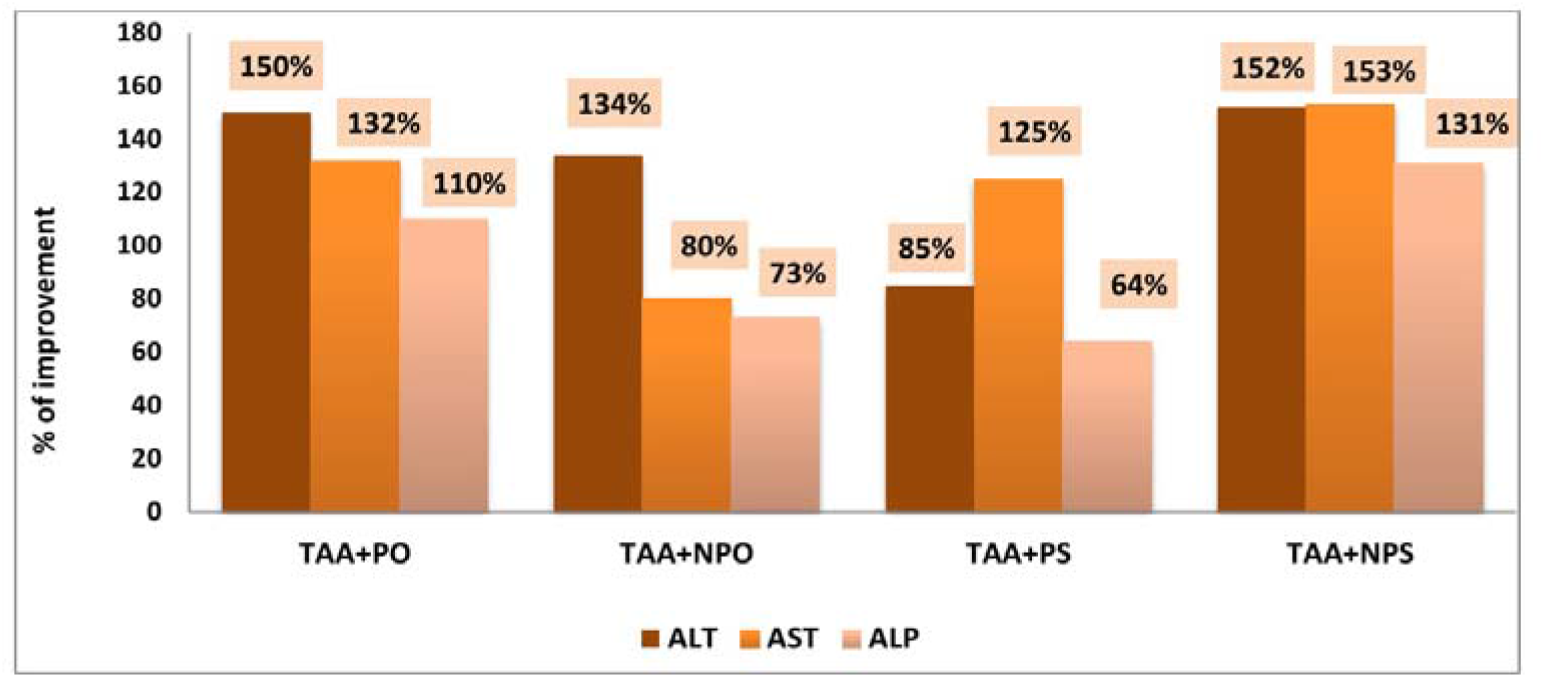
Therapeutic effects of both of Pomegranate and Pistachio and their nanoparticle forms on serum ALT, AST and ALP. oValues are expressed as % of improvement related to normal control and thioacetamide groups.

**Fig. 7:**
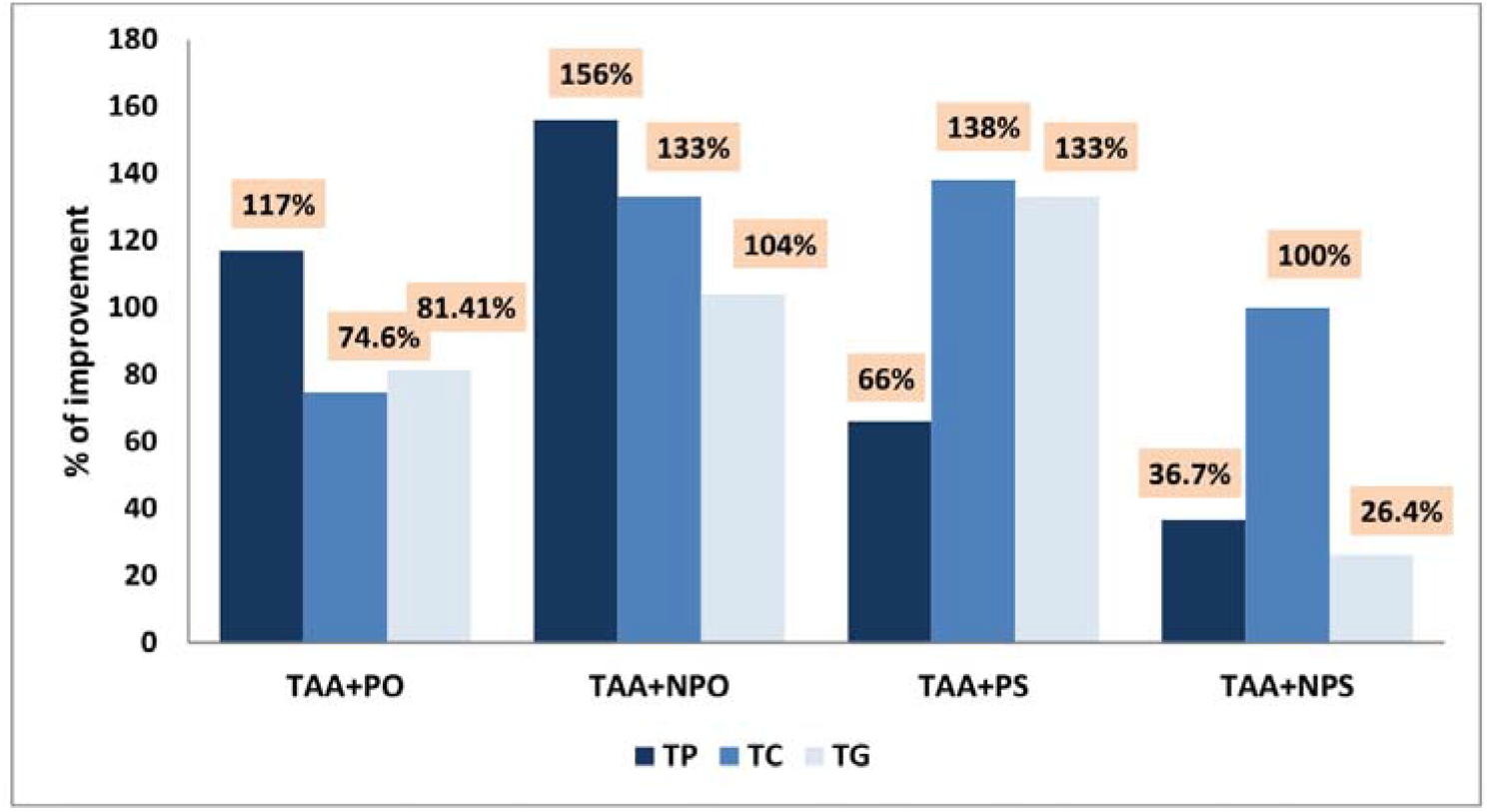
Therapeutic effects of both of Pomegranate and Pistachio and their nanoparticle forms on serum Total lipid, Total cholesterol and Total glyceride. *Values are expressed as % of improvement related to normal control and thioacetamide groups.

**Table. (2):** Therapeutic effects of both of *pomegranate, pistachio* and their nanoparticles on liver functions and lipid profile.

| Groups Parameters | Control (-ve) | TAA (+ve) | TAA+PO | TAA+NPO | TAA+PS | TAA+NPS |
| --- | --- | --- | --- | --- | --- | --- |
| ALT (U/ml) | 48.6<sup>c</sup><br>±<br>7.40 | 121.4<sup>a</sup><br>±<br>9.04 | 48<sup>c</sup><br>±<br>6.81 | 55.8<sup>c</sup><br>±<br>6.72 | 79.6<sup>b</sup><br>±<br>12.70 | 47<sup>c</sup><br>±<br>5.41 |
| AST (U/ml) | 54.4<sup>d</sup><br>±<br>6.58 | 157.2<sup>a</sup><br>±<br>10.70 | 85.25<sup>c</sup><br>±<br>7.95 | 113.8<sup>b</sup><br>±<br>9.67 | 89<sup>c</sup><br>±<br>4.94 | 74<sup>c</sup><br>±<br>17.70 |
| ALP (U/l) | 108.48<sup>cd</sup><br>±<br>17.16 | 233.90<sup>a</sup><br>±<br>15.72 | 124.42<sup>c</sup><br>±<br>14.90 | 154.28<sup>b</sup><br>±<br>16.58 | 164.48<sup>b</sup><br>±<br>24.27 | 91.42<sup>d</sup><br>±<br>5.60 |
| Total Lipid (TP) (g/dl) | 0.402<sup>c</sup><br>±<br>0.032 | 0.909<sup>a</sup><br>±<br>0.06 | 0.439<sup>c</sup><br>±<br>0.035 | 0.58<sup>b</sup><br>±<br>0.057 | 0.609<sup>b</sup><br>±<br>0.062 | 0.610<sup>b</sup><br>±<br>0.066 |
| Total cholesterol (TC) (mg/dl) | 40.20<sup>c</sup><br>±<br>1.92 | 90.02<sup>a</sup><br>±<br>2.12 | 63.49<sup>b</sup><br>±<br>2.14 | 36.22<sup>d</sup><br>±<br>2.18 | 34.52<sup>d</sup><br>±<br>0.71 | 36.05<sup>d</sup><br>±<br>2.70 |
| Total glyceride (TG) (mg/dl) | 96.32<sup>e</sup><br>±<br>12.51 | 297.1<sup>a</sup><br>±<br>18.9 | 146.08<sup>d</sup><br>±<br>15.12 | 196.6<sup>c</sup><br>±<br>22.45 | 278.2<sup>a</sup><br>±<br>9.73 | 250.8<sup>b</sup><br>±<br>8.75 |
| Total bilirubin (TB) (mg/dl) | 1.172<sup>d</sup><br>±<br>0.115 | 2.26<sup>b</sup><br>±<br>0.22 | 1.83<sup>c</sup><br>±<br>0.08 | 2.57<sup>b</sup><br>±<br>0.22 | 3.44<sup>a</sup><br>±<br>0.095 | 2.57<sup>b</sup><br>±<br>0.33 |

- Each group has a mean ± SD of ten rats. - Statistical analysis was conducted using a one-way variance analysis (ANOVA) using the Co-Stat computer program with the lowest level (LSD) between p<0.05 classes. - Groups with the same letters are not significantly different, whereas those with different letters vary considerably at p<0.05.

### 3.6. Effect ofpomegranate and pistachios in their normal and nanoforms on oxidative stress

GSH content, SOD activity, LPO and Nitric oxide (NO) levels were taken as markers of oxidative status. An obvious depletion was seen in the levels of GSH & SOD with a prominent increase in each of LPO and NO levels in the challenged TAA intoxicated groupat ***p***<0.0001 **(Table 3, Fig. 8.**).The treatment with Pomegranate and Pistachio (100 mg/kg) either in normal or nano forms effectively increased the levels of both GSH and SOD***p*** < 0.05. While levels of LPO and NO showed a prominent reduction upon treatment with both extracts **(Table 3, Fig. 8.)**. While its Nano form of pistachio revealed more powerful as it restored the NO value to 1.58 compared to 3.41 in the challenged TAA group. The TAA-intoxication resulted in a remarkable elevation at (***p*** < 0.05) in the measured inflammatory markers as 2 fold and 2.5 fold in IL6 and TNHα consecutively and about 6 fold increase in the levels of AFP as well as HSP_70_ compared to the TAA intoxicated group **(Table 3, Fig. 9.).** Treatment with either pomegranate peel and or pistachio normal form extracts markedly reduced AFP, TNFα, IL6 and HSP_70_ levelsrespectively with the advantage of pistachio either in normal or Nano forms as it revealed a higher restoration power than pomegranate with the inflammatory markers as seen in **Table 3**. Therefore, the results provided proof for each of the pomegranates and pistachios, either in their normal or nano forms, showing a different percentage of change as shown in Fig. 9

**Fig. 8.**
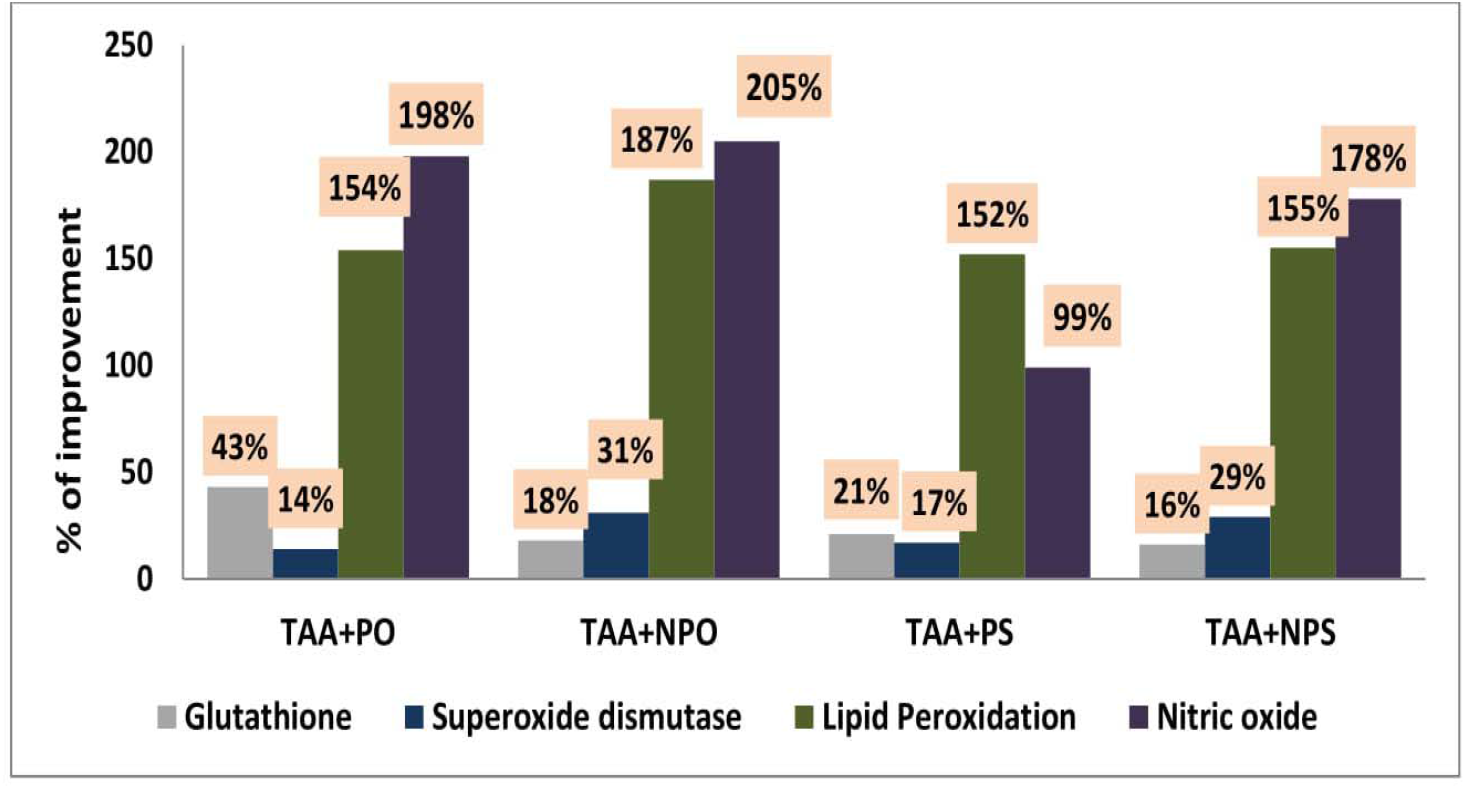
Therapeutic effects of both of Pomegranate and Pistachio and their nanoparticle forms on oxidative stress markers. *Values are expressed as % of improvement related to normal control and thioacetamide groups.

**Fig. 9.**
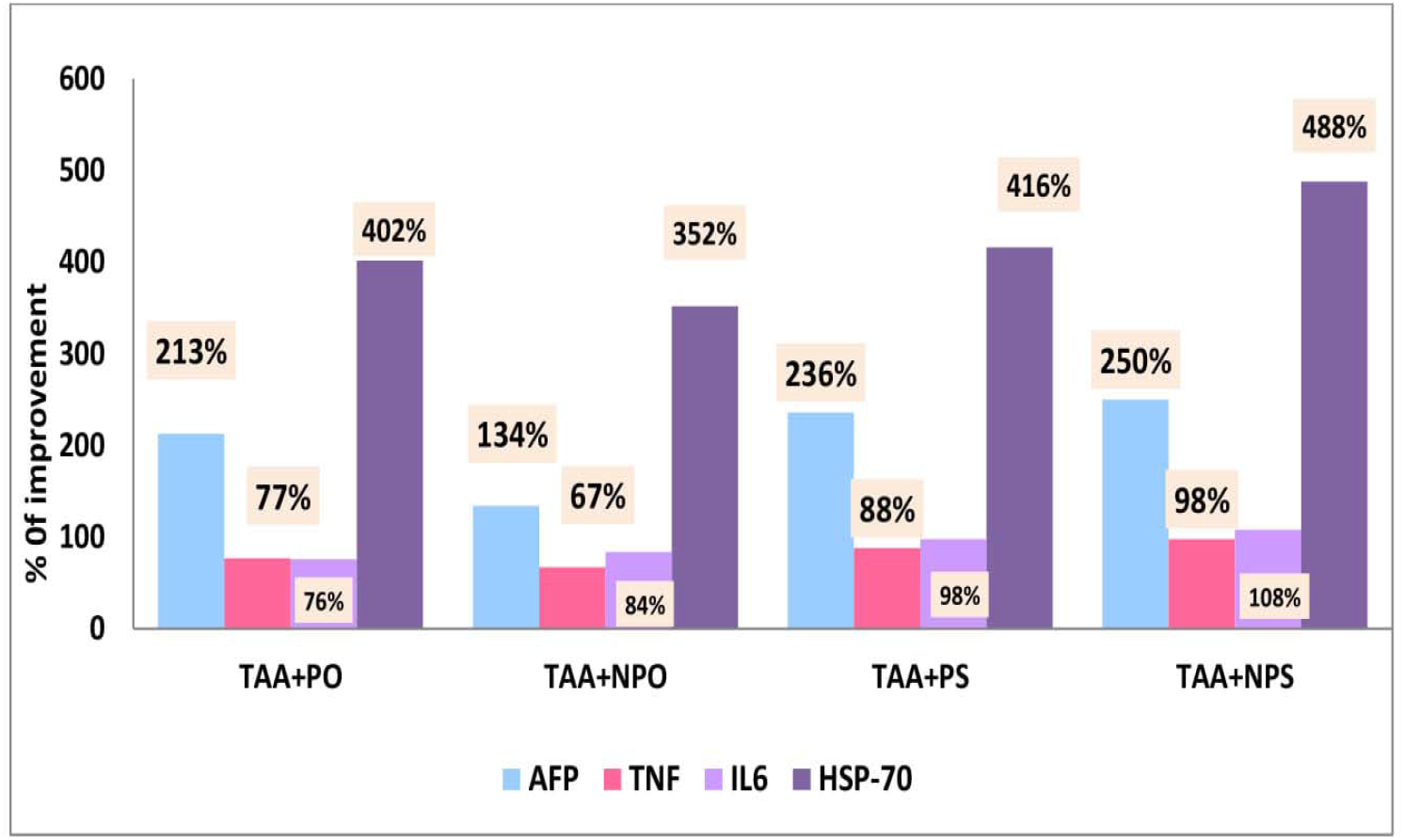
Therapeutic effects of both of Pomegranate and Pistachio and their nanoparticle forms on inflammatory cytokines. *Values are expressed as % of improvement related to normal control and thioacetamide groups.

**Table (3).** Therapeutic effects of both of pomegranate, pistachio and their nanoparticles on oxidative stress markers and inflammatory cytokines.

| Groups Parameters | Control (-ve) | TAA (+ve) | TAA+PO | TAA+NPO | TAA+PS | TAA+NPS |
| --- | --- | --- | --- | --- | --- | --- |
| **Glutathione (ug/mg protein)** | **48.50<sup>a</sup>**<br>±<br>2.62 | **21.80<sup>d</sup>**<br>±<br>3.02 | **42.20<sup>b</sup>**<br>±<br>2.40 | **30.43<sup>c</sup>**<br>±<br>1.68 | **31.98<sup>c</sup>**<br>±<br>3.06 | **29.97<sup>c</sup>**<br>±<br>4.13 |
| **Superoxide dismutase (Umol/mg protein)** | **0.123<sup>a</sup>**<br>±<br>0.008 | **0.048<sup>d</sup>**<br>±<br>0.004 | **0.065<sup>c</sup>**<br>±<br>0.007 | **0.087<sup>b</sup>**<br>±<br>0.009 | **0.069<sup>c</sup>**<br>±<br>0.006 | **0.083<sup>b</sup>**<br>±<br>0.006 |
| **Lipid Peroxidation (Umol /mg protein)** | **1.099<sup>d</sup>**<br>±<br>0.086 | **3.45<sup>a</sup>**<br>±<br>0.082 | **1.77<sup>b</sup>**<br>±<br>0.104 | **1.402<sup>c</sup>**<br>±<br>0.185 | **1.78<sup>b</sup>**<br>±<br>0.071 | **1.75<sup>b</sup>**<br>±<br>0.14 |
| **Nitric Oxide (mmol/g tissue)** | **1.034<sup>d</sup>**<br>±<br>0.18 | **3.41<sup>a</sup>**<br>±<br>0.31 | **1.37<sup>cd</sup>**<br>±<br>0.17 | **1.28<sup>cd</sup>**<br>±<br>0.12 | **2.39<sup>b</sup>**<br>±<br>0.14 | **1.58<sup>c</sup>**<br>±<br>0.21 |
| **AFP ( ng/ml)** | **1.212<sup>e</sup>**<br>±<br>0.07 | **7.88<sup>a</sup>**<br>±<br>0.27 | **5.304<sup>c</sup>**<br>±<br>0.19 | **6.26<sup>b</sup>**<br>±<br>0.058 | **5.03<sup>d</sup>**<br>±<br>0.13 | **4.85<sup>d</sup>**<br>±<br>0.135 |
| **TNF (Pg/ml)** | **60.58<sup>f</sup>**<br>±<br>0.89 | **159.90<sup>a</sup>**<br>±<br>2.14 | **112.54<sup>c</sup>**<br>±<br>0.855 | **119.7<sup>b</sup>**<br>±<br>0.86 | **106.82<sup>d</sup>**<br>±<br>1.61 | **100.94<sup>b</sup>**<br>±<br>2.08 |
| **IL6 (Pg/ml)** | **93.87<sup>f</sup>**<br>±<br>2.39 | **236.77<sup>a</sup>**<br>±<br>9.95 | **165.95<sup>b</sup>**<br>±<br>2.71 | **158.36<sup>c</sup>**<br>±<br>1.89 | **145.67<sup>d</sup>**<br>±<br>3.60 | **135.47<sup>e</sup>**<br>±<br>2.39 |
| **HSP-70 (PG/mg)** | **111.75<sup>f</sup>**<br>±<br>4.90 | **791.69<sup>a</sup>**<br>±<br>6.65 | **344.24<sup>c</sup>**<br>±<br>4.95 | **400.34<sup>b</sup>**<br>±<br>7.40 | **329.39<sup>d</sup>**<br>±<br>6.56 | **249.78<sup>e</sup>**<br>±<br>4.74 |

The data are represented as average ± SD of 10rats in each group. Statistical analysis was conducted using a one-way variance analysis (ANOVA) using the CoStat computer program with the lowest level (LSD) between p<0.05 classes. • Groups with the same letters are not significantly different, whereas those with different letters vary considerably at p<0.05.

### 3.7. *Effect of pomegranate and pistachio* in their normal and Nano forms onDNA degradation

Comet assay is a sensitive and well-validated technique used to evaluate the fragmentation of DNA, a typical component of toxic DNA damage and apoptosis. Compared to the normal control group, TAA has substantially caused DNA damage. This DNA damage was demonstrated by a significant increase in the comet parameters presented as tail length with different three classes **(Fig. 10, 11)**, Tail and tail moment DNA (percentage); the primary predictive parameter for DNA damage **(Table 4).** In the hepatotoxic community there was a large broom-like tail and a small comet eye.**(Fig. 10, 11)**. The length of the tail increases as DNA damage increases as seen in **Fig.10, 11.** Whereas, the administration of both pomegranate and pistachio in either their natural or nano forms impaired the TAA effect by substantially reducing the length of the tail; with the dominance of the pomegranate and pistachio nano forms (Table 4, Fig. 12).

**Fig. 10.**
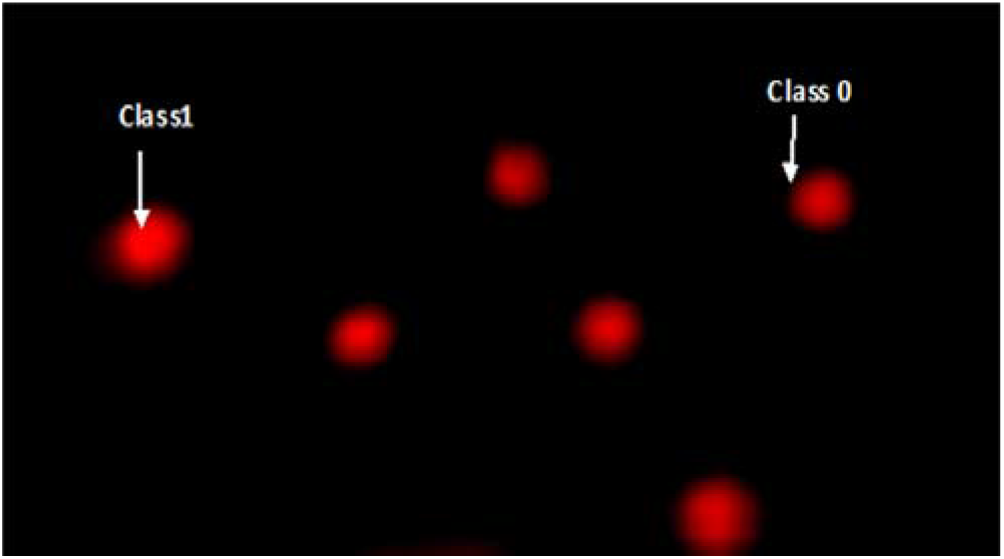
Visual DNA damage score (class 0 and 1) using comet assay in liver tissues.

**Fig. 11.**
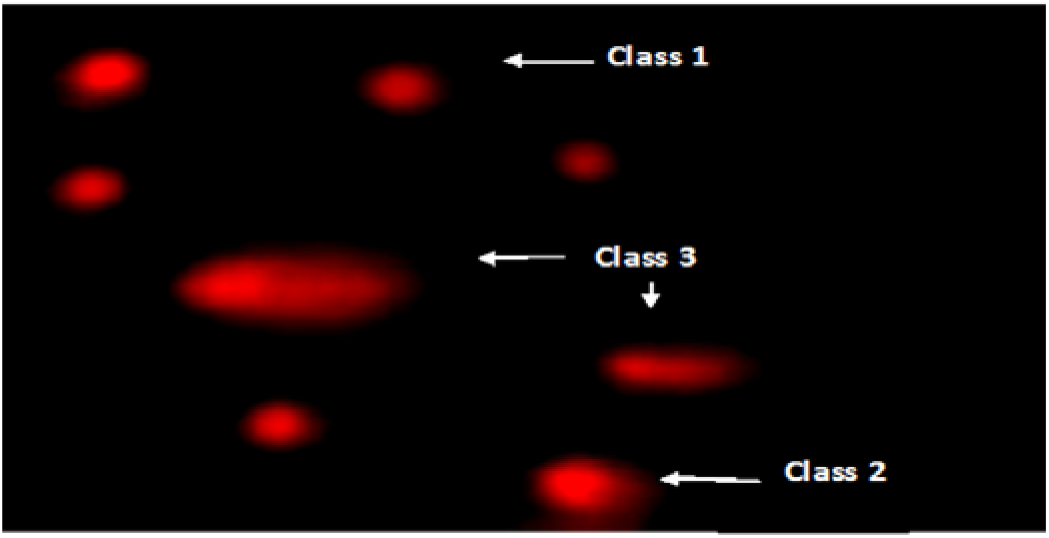
Visual DNA damage score (**classes 2 and 3**) using liver tissue comet assay.

**Fig. 12:**
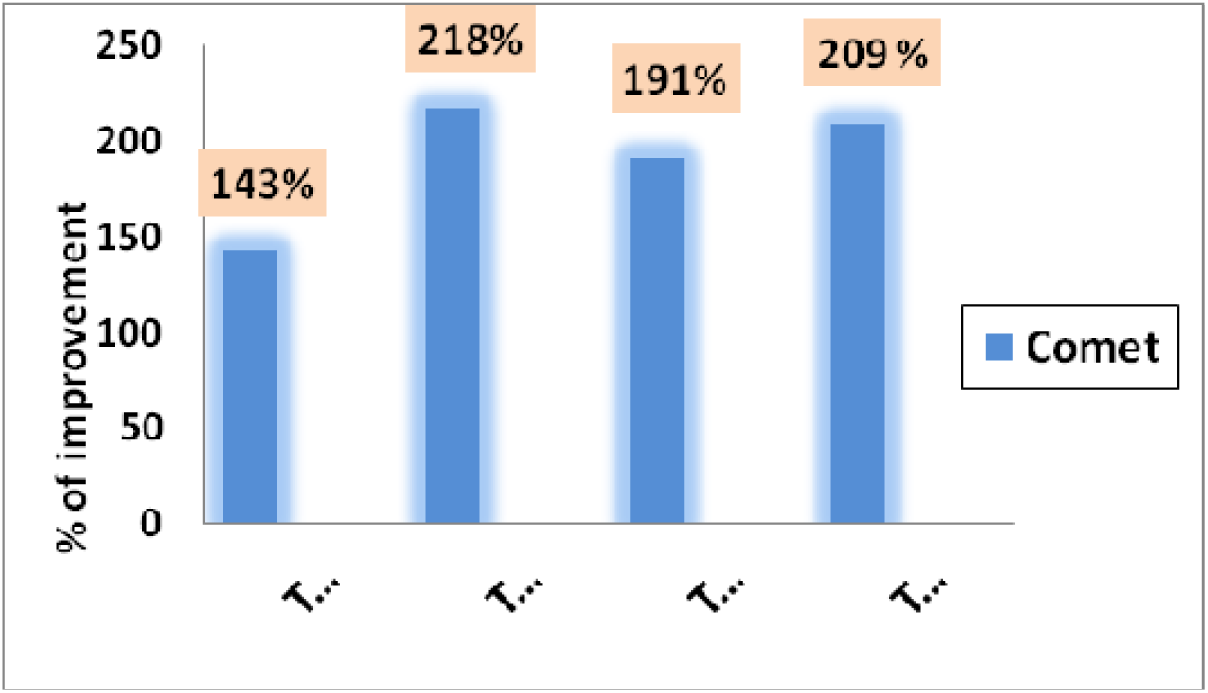
Therapeutic effects of both of Pomegranate and Pistachio and their nanoparticles forms on Comet assay. *Values are expressed as % of improvement related to normal control and thioacetamide groups.

**Table (4):** Therapeutic effects of both of pomegranate and pistachio and their nanoparticles on DNA damaged cells by comet assay.

| Groups | cells No. | | Comet class | | | | Damaged DNA cells<br>(mean ± SD) |
| --- | --- | --- | --- | --- | --- | --- | --- |
|  | Analyzed | Total comets | 0 | 1 | 2 | 3 |  |
| Negative control | 500 | 33 | 467 | 25 | 8 | 0 | 6.61 ± 0.54<sup>d</sup> |
| TAA | 500 | 123 | 377 | 35 | 43 | 45 | 24.64 ± 1.14<sup>a</sup> |
| TAA+PO | 500 | 73 | 427 | 28 | 26 | 19 | 14.62 ± 0.89<sup>b</sup> |
| TAA+NPO | 500 | 51 | 449 | 24 | 17 | 10 | 10.20 ± 1.3<sup>c</sup> |
| TAA+PS | 500 | 61 | 439 | 21 | 23 | 17 | 12.22 ± 1.4<sup>bc</sup> |
| TAA+NPS | 500 | 54 | 446 | 16 | 22 | 16 | 10.81 ± 0.8<sup>c</sup> |

Class 0= no tail; 1= tail length < diameter of nucleus; 2= tail length between 1X and 2X the diameter of nucleus; and 3= tail length > 2X the diameter of nucleus. (\*): No of cells analyzed was 100 per an animal.

### 3.8. Pearson’s correlations and ROC-Curve of the parameters in all groups

**Table 5** shows positivecorrelations between liver functions, lipid profile, oxidative stress markers and inflammatory cytokines at different correlation levels expect for GSH and SOD activity.**Table 6** shows area under the curve AUC equal 1 and showed the same sensitivity and specificity values for virtually every biomarker.

**Table (5).** Pearson’s correlations between liver function, lipid profile, oxidative markers and inflammatory cytokines.

| **R**<br>**(Person Correlation)** | **AFP** | **TNFα** | **IL6** | **HSP70** |
| --- | --- | --- | --- | --- |
| **ALT** | 0.621** | 0.735** | 0.773** | 0.849** |
| **AST** | 0.857** | 0.905** | 0.889** | 0.928** |
| **ALP** | 0.694** | 0.778** | 0.801** | 0.863** |
| **TG** | 0.733** | 0.713** | 0.643** | 0.653** |
| **TC** | 0.571** | 0.714** | 0.835** | 0.816** |
| **TP** | 0.763** | 0.821** | 0.805** | 0.858** |
| **TB** | 0.513** | 0.398** | 0.263 | 0.244 |
| **GSH** | -0.804** | -0.786** | -0.684** | -0.734** |
| **SOD** | -0.854** | -0.885** | -0.873** | -0.830** |
| **MDA** | 0.759** | 0.862** | 0.916** | 0.924** |
| **NO** | 0.680** | 0.768** | 0.804** | 0.832** |
| **AFP** | 1 | 0.977** | 0.979** | 0.886** |
| **TNFα** | 0.977** | 1 | 0.773** | 0.849** |
| **IL6** | 0.921** | 0.979** | 1 | 0.928** |
| **HSP70** | 0.886** | 0.961** | 0.979** | 1 |

\* The Correlation is significant at the level of 0.05. \* \* The correlation is significant at the level of 0.01.

**Table 6:**
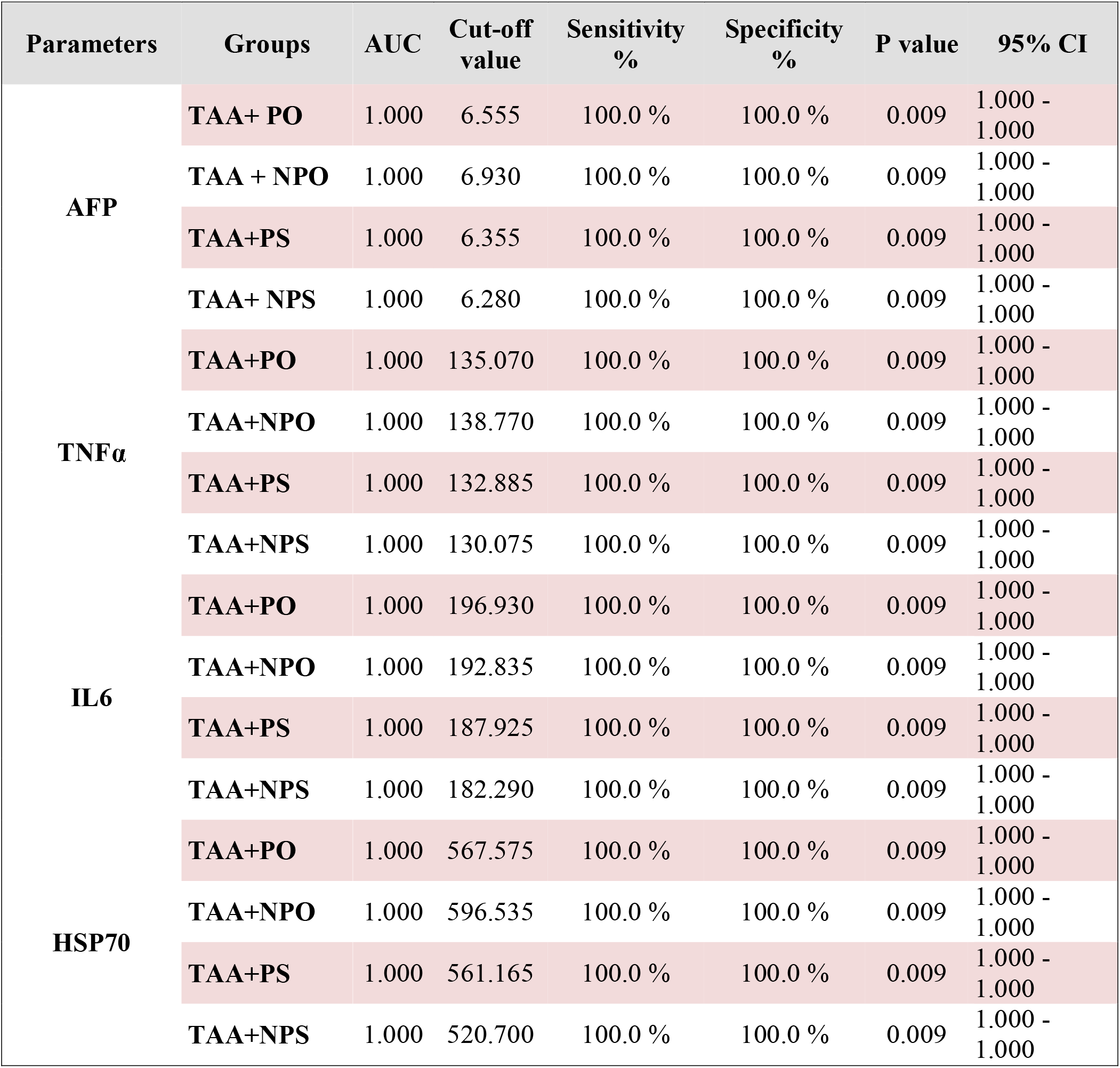
Sensitivity & specificity of the measured parameters in all groups (ROC-curve)

### 3.9. Histopathologic examination

Histopathological analysis of the representative liver tissues of the control group showed normal morphology, hepatocyte structure and central vein. **(Fig. 13.A)**. TAA-intoxicated group Displayed Focal inflammatory cell infiltration in the hepatic parenchyma covering the central dilated vein **Fig. 13. B,** with diffuse spread of kupfer cells between hepatocytes. Pomegranate and /pistachio treatment (100 mg/kg) either in normal or nano forms have shown observable healing effect presented as shown in **(Fig. 13.C, D, E, F)** with only some degenerative changes.

**Fig. 13.**
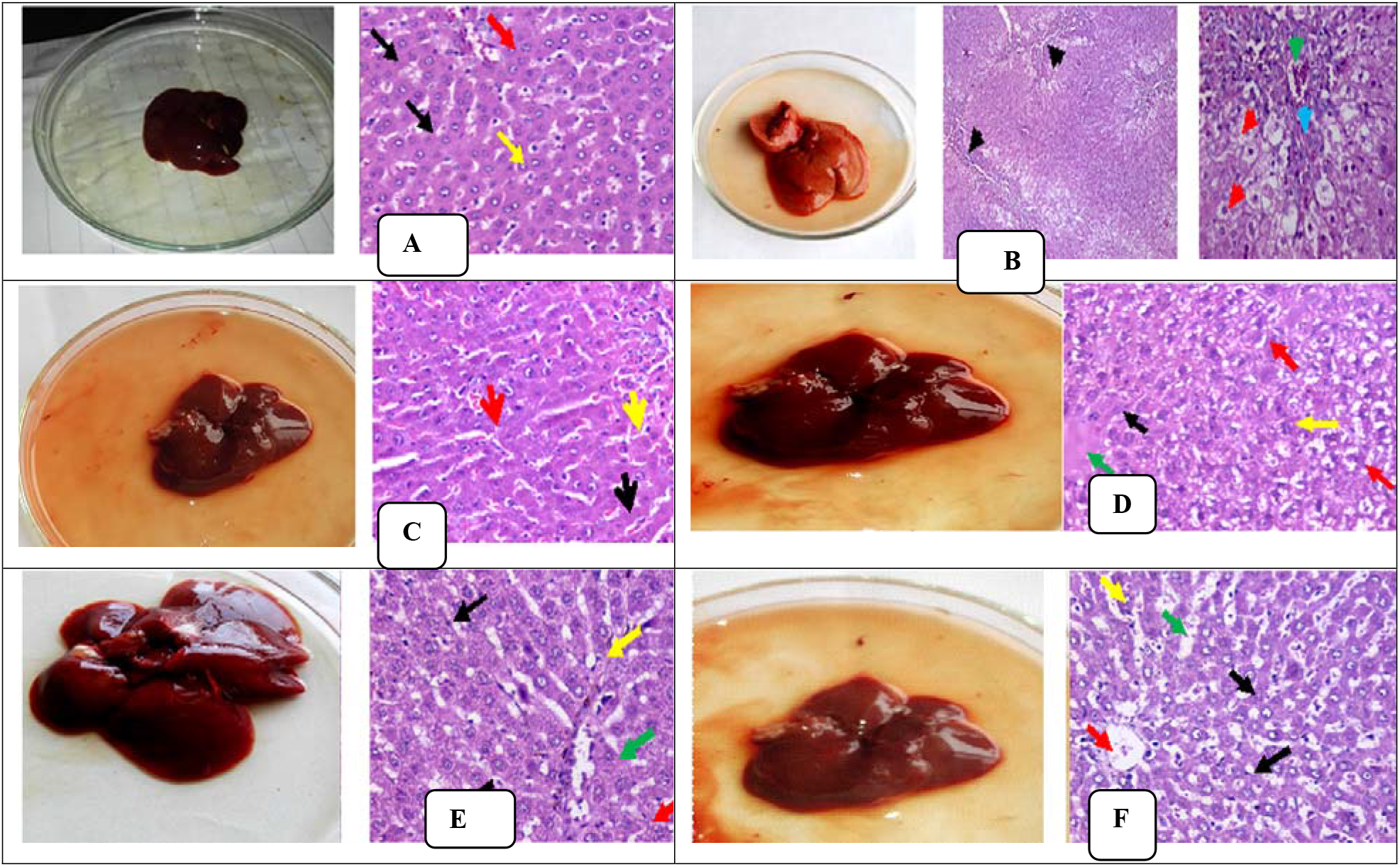
**A.** The liver portion of the normal control group (standard control) showed typical structure and morphology of the liver tissue, thin-plated hepatocytes (black arrow) and sinusoids (yellow arrow), central red vein. (H&E, x200, x400). **B.** Liver section from animal with TTA group showed partial distortion of lobular hepatic architecture (black arrow), with moderate hydropic degeneration of hepatocyte (red arrow) and binucleated hepatocytes (yellow arrow), portal tract with vein congestion (green arrow) thick fibrous tissue in portal tract (blue arrow) (H&E,x200, x40). **C.** liver section of animal treated with pomegranate group displayed Hepatocytes arranged in thin plates (black arrow) and congested central vein (red arrow) with a preserved architecture, moderately dilated and congested sinusoids (yellow arrow) (H&E, x200, x400). **D.** Liver section of animal treated Nano pomegranate group showed intact lobular hepatic architecture, hepatocyte with thin plates (black arrow). Moderate hydropic degeneration (red arrow) and binucleated hepatocytes (yellow arrow), central vein (green arrow) (H&E, x200, x400). **E.** The liver section of the animal treated with Pistachio extract showed hepatic tissue intact lobular hepatic structure and architecture, hepatocytes arranged in thin plates with mild hydropic changes (black arrow) and congested central vein (red arrow) binucleated hepatocytes (yellow arrow), slightly dilated and congested sinusoids(green arrow) (H&E, x200, x400). **F.** liver section of animal treated with Nano Pistachio group showed Hepatic tissue with nearly normal structure and architecture, hepatocytes arranged in thin plates (black arrow) and congested central vein (red arrow) binucleated hepatocytes (yellow arrow), mildly dilated and congested sinusoids (green arrow) (H&E, x200, x400).

### 3.10 Effect of pomegranate / pistachio either in their normal and/or their Nano forms on the inflammatory markers by immunohistochemistry

The anti-inflammatory effect of pomegranate /pistachio either in normal and/or its nano forms was further evaluated viaimmunohistochemistry expression of pro inflammatory markers, IL-6 TNHx in hepatic tissues **(Fig. 14 A - L)**. TAA intoxication 350 mg/kg resulted in a substantial elevation in IL-6 and TNF-α expression compared to normal control group **(Fig. 14, A, G)** respectively that was located mostly in cytoplasm of hepatocytes in portal tract and fibrous septa. Treatment with pomegranate/and or pistachio either in their normal or Nano forms improved and down regulated the expression of both IL6 (**Fig.14, B-F)** and TNF-**α (Fig.14, I - L)** respectively, with the notion has been shown that pomegranate /pistachio in its nano forms is more likely to inhibit TAA induced inflammatory response compared to positive control **(Fig.14. B - L).**

**Fig. 14:**
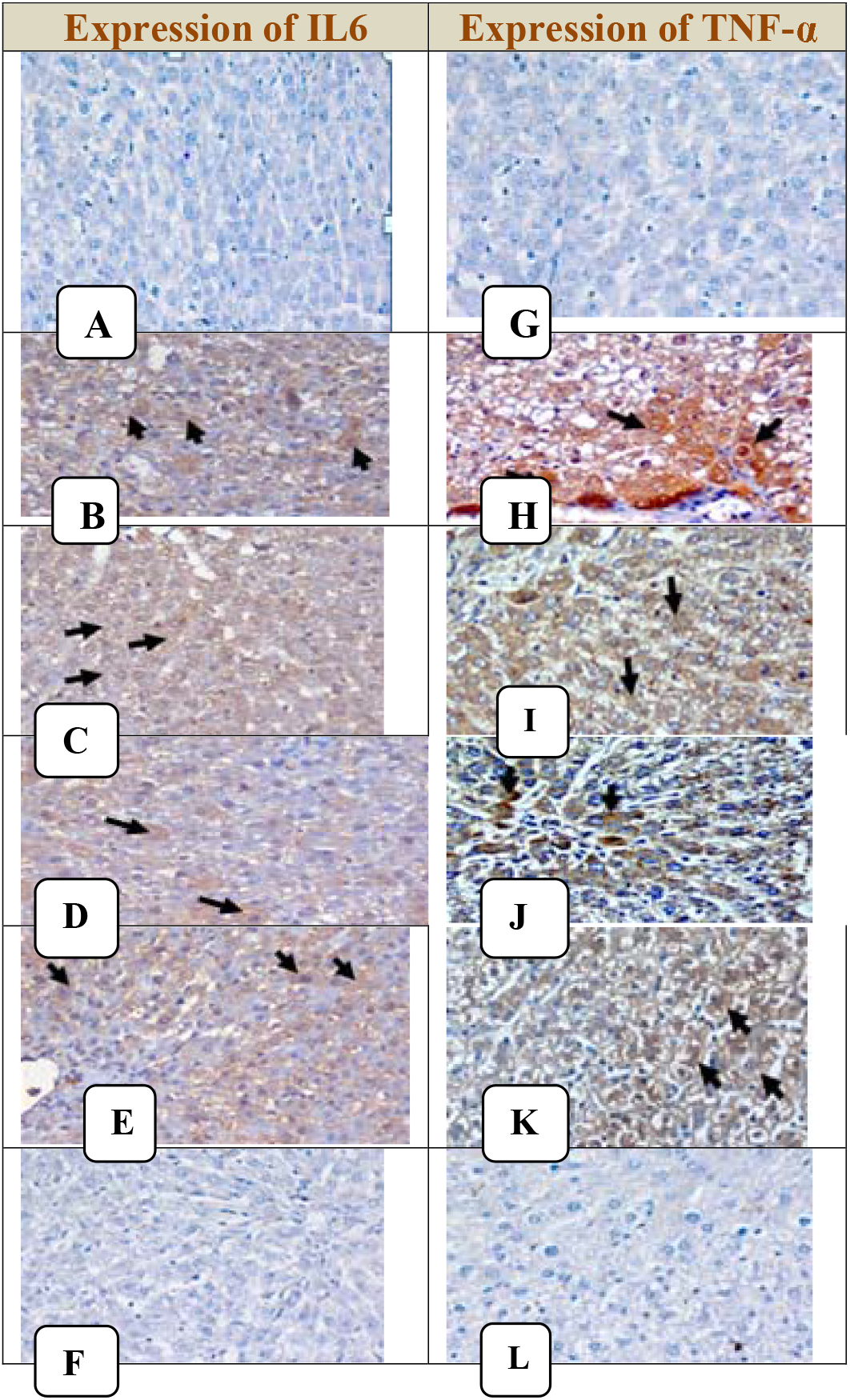
**A.** liver section from normal control group showed negative expression of IL6 (DAB,IHC, x400). **B.** liver section from positive control group showed moderate expression of IL6 as brownish cytoplasmic stain of hepatocytes in about 60% (black arrow)( DAB,IHC, x400). **C.** liver section from animal treated with pomegranate group showed moderate expression of IL6 as brownish cytoplasmic stain of hepatocytes in about 40% (black arrow)( DAB,IHC, x400).**D.** liver section from animal treated Nano pomegranate group showed moderate expression of IL6 as brownish cytoplasmic stain of hepatocytes in about 10% (black arrow)( DAB,IHC, x400).**E.** liver section from animal treated Pistachio group showed mild expression of IL6 as brownish cytoplasmic stain of hepatocytes in about 25% (black arrow)( DAB,IHC, x400). **F.** liver section from animal treated Nano Pistachio group showed negative expression of IL6 (DAB, IHC, x400). **G.** liver section from normal control group showed negative expression of TNFα (DAB,IHC, x400).**H.**liver section from positive control group showed marked expression of TNF ***α*** as brownish cytoplasmic stain of hepatocytes in about 70% (black arrow)( DAB,IHC, x400).**I.**liver section from animal treated with pomegranate group showed moderate expression of TNF ***α*** as brownish cytoplasmic stain of hepatocytes in about 40% (black arrow)( DAB,IHC, x400).**J.**liver section from animal treated Nano pomegranate group showed moderate expression of TNF ***α*** as brownish cytoplasmic stain of hepatocytes in about 10% (black arrow)( DAB,IHC, x400).**K.**liver section from animal treated Pistachio group showed moderate expression of TNF α as brownish cytoplasmic stain of hepatocytes in about 30% (black arrow)( DAB,IHC, x400).**L.**liver section from animal treated Nano Pistachio group showed negative expression of TNF α (DAB, IHC, x400).

## 4. Discussion

Oxidative stress appears to play a major role in the outbreak and progression of wide array of diseases, including the liver^[49]^. TAA has been recognized as hepatotoxic compound in different studies^[50]^. As it transforms into thioacetamide sulfoxide (TAAS) along the pathway-dependent cytochrome P-450 (CYP) and/or flavin-containing monooxygenase systems forming TAA sulfoxide involving TAA-S, S-dioxide that begin to form lipid peroxidation at the plasma membrane level^[51, 52]^. Binding of these metabolites to liver macromolecules dramatically inspire a reactive oxygen species moiety which in turn induce an acute centrilobular liver necrosis and consequently leads to liver diseases and further can overwhelm the body antioxidant defense mechanisms reaching to vital ingredients ending up to damage the DNA^[12]^. Emerging research evidence indicates that natural compounds / plants have shown beneficial effects of antioxidants in the defense of the liver against TAA-induced injury^[53]^ and can improve immune function^[54]^ either in normal or in nano-form^[55]^.

In accordance with this assumption,the results of this study demonstrated the efficacy of the nanoparticles synthesized from the natural product as an excellent capping and reducing agent capable of producing pomegranate and pistachio nano particles [***P. GranatumL***] [***pistashiaatlantica***] aqueous nano-extracts with average spherical size ranging from 7-17 nm. The present study showed that TAA-intoxication in a single dose 350 mg/bw had superabundant hepatotoxic symptoms clearly represented via the elevation in liver enzymes (AST, ALT and ALP), lipid profiles (TP, TC and TG) as well as total bilirubin **(Table.2)**. This observation was close to Fontana et al^[56]^ and Al shawsh et al^[57]^ findings as they demonstrated that TAA has been shown to interfere with the continuous movement of RNA from the nucleus to the cytoplasm, may cause membrane injury resulting in elevation and release of serum markers of the liver^[58]^. Such leakage of the enzymes into the blood was accompanied in our study with partial distortion of lobular hepatic architecture, with moderate hydropic degeneration of hepatocyte and binucleated hepatocytes, portal tract with vein congestion thick fibrous tissue in portal tract in addition to inflammatory infiltration of the liver which revealed severe liver injury **(Table 2, Fig 13B)**. Interestingly, our findings demonstrated an obvious amelioration and improvement in all parameters when treated with pomegranate peel extract and /or pistachio either in normal or nano forms as indicated by reduction or normalization of the levels of aminotransferases, alkaline phosphatase, lipid profile, total bilirubin **(Table 2, Fig. 6, 7).** These improvements extends to liver architecture (**Fig 13C-F**). These observations are most likely attributed to the presence of hydrolysable polyphenols, specifically ellagitannins, which were considered to be the most active antioxidants among the tannins contained therein. These compounds (elagic acid, punicalagin, punicalin and galagid acid) have been shown to have increased antioxidant and pleiotropic effects and are believed to act in synergy^[59, 60]^ Our findings were found to be similar to studies of Husain et al^[61]^, Bachoual et al^[62]^ and seem to be compatible with Esmaillzadeh et al^[63]^ as they demonstrated that pomegranate juice significantly reduce the total cholesterol level. The results also go in consistent with Al-Shaaibi et al^[64]^ as they mentioned that administration of Pomegranate peel extract ameliorate liver enzymes, control lipogenesis and improve the histologic image of liver. And in agreement with Kocyigit et al^[65]^, Sheridan et al^[66]^; Kay et al^[67]^as they demonstrated that pistachio has a beneficial effect on the improvementof aminotransferase levels and blood lipid profile, This can be attributed to the existence of ellagitannins hydrolysable polyphenols as a highly antioxidant compounds, similar to that in pomegranate.In addition to other antioxidants that can effectively play a key role in stopping oxidative reactions in chains such as fatty acids, flavonoids and phenolic compounds.^[32, 33]^ tocopherols, carotenes, lutein, selenium and phytoestrogens^[68, 69]^.

In another observation, both nano forms of pomo and pistachio treated groups showed significant increase levels of bilirubin. The increased levels was further describes nano forms of pistachio as a protective polyphenols. These findings appear to be consistent with previous data related to Bilirubin IXα as it recognized as a potent antioxidant by Vitek et al^[70]^. While bilirubin is toxic in children, it is considered for health promotion in adults. At the same time, according to some clinical literature, moderately elevated serum bilirubin levels were generally correlated with decreased incidence of chronic diseases^[71]^.

A pronounced increment in lipid peroxidation and NO levels were recorded in hepatic homogenate. These results were concomitant with a pronounced, significant reduction in hepatic - GSH and SOD levels. The results can be delegated to the thiono-sulfur compound of TAA as a hepatotoxic which induce hepatic damage and act as an initiator for a series of severe inflammatory changes via generation of ROS and down regulator of the antioxidant defense mechanism^[72]^. Moreover In liver disease L-arginine-NO pathway is activated, and lead to the formation of nitric oxide, the mater which can explain the significant increase in nitric oxide level upon intoxication by TAA^[73]^ **(Table 3).**

Interestingly, the great significant reduction effect on the lipid peroxidation level as well as nitric oxide synchronized with increment in glutathione and SOD levels were taken by the nano forms of each of pomegranate & pistachio extracts respectively as they proved to be more superior compared to their normal form **(Table 3, Fig. 8)**. The improvement registered in the measured redox parameters may be in agreement with what previously reported that Stimulation of the Nrf2/ARE pathway is necessary for antioxidant defense enzyme induction and intracellular GSH modulation in response to stress^[74]^ **(Table 3)**. These nano form results can be interpreted upon their Benign nature and the presence of essential functional groups make AgNPs very stable and prevent nanoparticles from being anchored.In addition, the existence of the bioactive molecules on the surface of AgNPs may increase its efficacy and bioavailability. It can also enhance its solubility, improve plasma half-life, preventing degradation in the intestinal environment and increasing permeation in the small intestine which may confer strong antioxidative activity^[75]^.

Our findings are further consistent with the previous study that pomegranate peel extract had the highest free radical scavenging capacity among the medicinal plants tested and not only juice and oil seed^[76, 77]^. And another study for Alturfan et al^[78]^ demonstrated that pistachio consumption can provide antioxidant protection.

In the current study and following treatment with TAA high plasma levels of AFP, TNF-α, IL6, and HSP70 were observed in serum as well as in tissue (**Table 3, fig. 9**). These results may really meet the fact that alteration in the redox status would stimulate the entire inflammatory cascade including the activation of a series of transcription factors^[79, 80, 81]^. AFP is regarded as a biomarker of proliferating liver stem cells under conditions of liver injury^[82]^.Our data showed a marked increase in AFP’s post-TAA liver content by about 6 fold compared to [1.2ng/ml] in normal group **(Table 3)**, nano pistachio was in front of its normal and other pomo forms to attenuate and improve the AFP over-production and protect its normal transcription levels in liver, This may explain the mechanism by which nano pistachio worked to prevent its abnormal expression by binding to the enhancer region of the AFP gene. It also may encourage the fetal hepatocyte to start to regenerate. This observation is in accordance with Hessin et al^[83]^ who demonstrated that Pistachio has the ability to modulate the alpha fetoprotein transcriptional levels.

TNF-α is a community of cytotoxic pro-inflammatory cytokines, believed to play a vital role in liver damage initiation^[84]^. Increased circulating TNF-α activates cell surface TNF-α receptors, resulting in activation of the stress-related protein kinases JNK and IKKβ, leading to increased development of additional inflammatory cytokines such as IL6. TNF reduction is considered a treatment to reduce damage to the liver^[85]^. Surprisingly attenuation of the TAA-induced up regulation of the cytokines TNFα, IL6 together was achieved by pomegranate and pistachio either in normal or in nano forms. These data seem to be parallel with coursodonboydle^[86]^who reported that increased levels of proinflammatory cytokine IL 6 in rat ileum model necrotizing enterocolitis have been normalized after POM seed oil treatment and also go in accordance with the reports of Grace et al^[87]^and Paternitiet al^[88]^, where the first revealed that Pistachio extract significantly reduced the non-mitochondrial oxidative burst in macrophages associated with inflammatory response.While the other attributed the decrease in TNF-α and Interleukins production to the presence of polyphenols in pistachios which possess antioxidant and antiinflammatory properties. These data were demonstrated immunohistochemicaly in **Fig.14, AL.**

On the other hand Heat-shock proteins (HSPs) are unique proteins that are synthesized rapidly in cells in response to different kinds of stress, and acting as intracellular protein repair^[89]^. A marked increase in HSP70 level was recorded following the single injection of TAA to be online with Kuramochi et al^[90]^ who demonstrated that mRNA expressions of HSP 70 tended to be increased with injection of TAA. However, in our study down regulation of HSP70 expression level was achieved by each extracts (Pomo and Pistachio) by different percentages **(Table 3, fig.9)**, but again pistachio extract in its nano form was the most powerful as an anti-inflammatory compared to pomegranate. This is may be interpreted upon the powerful flavonoids beside other phenolic compounds benzoic acid, and hydroxycinnamic acid derivatives, that have antioxidant properties exist in the nano pistachio to induce strong HSP70 immunoreactivity, prevent protein denaturation and succeeded to overcome TAA induced liver damage^[91, 92]^. These data are parallel to what previously reported by Ibrahim et al.^[93]^ who demonstrated that POM has the ability to prevent protein denaturation, as a way to combat hepatic oxidative stress caused by CCl4 and also when he attributed the suppression of heat shock protein 70 and inducible nitric oxide synthase 3-nitrotyrosine to pomegranate emulsions^[94]^.

Moreover, the current data demonstrated an excessive amount of ROS after surpassing various endogenous anti-oxidative defensive mechanisms oxidized the DNA biomolecules **(Fig10, 11).** Our finding was in agreement with Afifi et al^[95]^,Zargar et al^[15]^who found a rise in hepatocyte DNA fragmentation following injection of TAA which may be attributed to the increase in cell death via both apoptosis and necrosis^[96]^. However treatment with pomegranate and /or pistachio either in normal or Nano forms as antioxidants were significantly improved and increased DNA resistance to oxidant damage with the superiority of the Nano forms of both plant extracts **(Table 4, fig.12)**. The current results may be in consistence with that study of Ahmed et al^[97]^as he revealed that pomegranate extract with its strong antioxidants, antiinflammatory, anti-apoptotic and ATP-replenishing was able to protect rats from DNA damage. Same data was confirmed by Majidinia et al^[98]^ as he proved polyphenols rich green extracts which is the main constituent of Pistachio leaves and Pomegranate peel as the main effector in decreasing the ROS level and DNA fragmentation, Eventually, the authors verified the findings by positive correlations between the entire markers measured **(Table 5)** Specificity & sensitivity for all markers were further determined by Roc curve and the region under the curve was shown to be equal to 1 **(Table 6)** which concluded that the measured parameters are good markers in case of acute liver failure.

## 5. Conclusion

Our findings indicated the potential and beneficial role of either pomegranate or pistachio in their normal forms in general and/or in their nano particles in particular, improving the deleterious effects of TAA oxidative stress-mediated damage and reinforcing the antioxidant defense mechanism through its rich antioxidant and anti-inflammatory properties material. Moreover it also reflects the potential of plant extracts to integrate biomolecules that serve as both a reduction and stabilization agents for producing stable and shape-controlled AgNPs with effective medicinal benefits that could contribute to the development of new drugs for acute liver disease.

## Acknowledgement

Authors gratefully acknowledge the assistance and any academic help they provided during this study.

